# Attenuating payload potency expands the therapeutic window of immunotoxins

**DOI:** 10.64898/2026.09.27.754776

**Authors:** Huazhu Liang, Greg L. Beilhartz, Xindi Wang, Kangni Zou, Shi Bo Cao, Roman A. Melnyk

## Abstract

Immunotoxins (ITs) are potent targeted cancer therapeutics that couple tumor-selective receptor binding with efficient intracellular delivery and cytotoxic enzymatic activity of bacterial toxins. While ITs have shown impressive efficacy in hematologic malignancies, dose-limiting toxicity remains a major barrier to their broader clinical application, particularly in solid tumors. The exceptional potency of immunotoxin payloads contributes to off-tumor toxicity because even limited uptake by receptor-low cells can be sufficient to cause cell death. Here, we systematically investigated how receptor-binding domain (RBD) affinity and avidity interact with catalytic payload activity to determine IT selectivity. Using CD123-, HER3-, and HER2-targeted ITs, we found that increasing receptor-binding affinity or avidity altered potency but provided limited control over the selectivity between receptor-high and receptor-low cells. In contrast, attenuating catalytic activity produced profound gains in selectivity when payload potency was appropriately matched to receptor engagement. The optimal level of attenuation varied with receptor context and toxin scaffold, demonstrating that maximal selectivity emerges from balancing cellular delivery with catalytic activity rather than maximizing either property alone. *In vivo*, payload attenuation increased the maximum tolerated dose of a HER2-targeted IT by 400-fold and enabled robust antitumor activity well below its toxicity limit, whereas the WT counterpart showed no detectable efficacy at tolerated doses. Together, these findings uncover and establish payload activity as a tunable determinant of immunotoxin selectivity and provide a framework for matching receptor engagement with catalytic potency to develop safer and more broadly applicable targeted toxins.

## Introduction

Cancer cells frequently gain a fitness advantage by overexpressing cell-surface receptors that promote their growth and survival. This same receptor overexpression, however, creates a therapeutic vulnerability by distinguishing cancer cells from normal cells. For example, HER family receptor tyrosine kinases, such as HER2, are overexpressed in multiple tumor types, where they drive cell proliferation and survival^1–5^. These cancer-associated receptors can therefore be exploited to preferentially target therapeutics to malignant cells^6^.

Differential receptor expression has been widely exploited to selectively deliver potent cytotoxic payloads to cancer cells, most notably through antibody-drug conjugates (ADCs) and immunotoxins (ITs). Whereas ADCs deliver small-molecule cytotoxins, ITs harness bacterial toxin-derived catalytic and translocation domains to deliver an exceptionally potent cytotoxic enzyme into the cytosol. ITs are most commonly derived from protein toxins such as diphtheria toxin (DT; Fig. 1A) and *Pseudomonas* exotoxin A (PE)^7,8^, and consist of a highly potent cytotoxic enzyme domain linked to a targeting moiety that binds to cancer-associated surface receptors. Following receptor binding and internalization, the catalytic payload is translocated into the cytosol, where it disrupts essential cellular functions. In DT and PE, the catalytic domains are ADP-ribosyltransferases that irreversibly modify elongation factor-2 (EF-2), arresting protein synthesis and ultimately causing cell death (**Fig. 1A**). To date, three ITs have been approved for hematological malignancies: denileukin diftitox (Ontak^®^), a DT-based IL-2Rα/CD25 targeted-IT for cutaneous T-cell lymphoma^9^, tagraxofusp (Elzonris^®^), a DT-based IL-3Rα/CD123-targeted IT for blastic plasmacytoid-cell neoplasms^10^, and moxetumomab pasudotox (Lumoxiti^®^), a PE-based CD22-targeted IT^11^.

**Figure 1.**
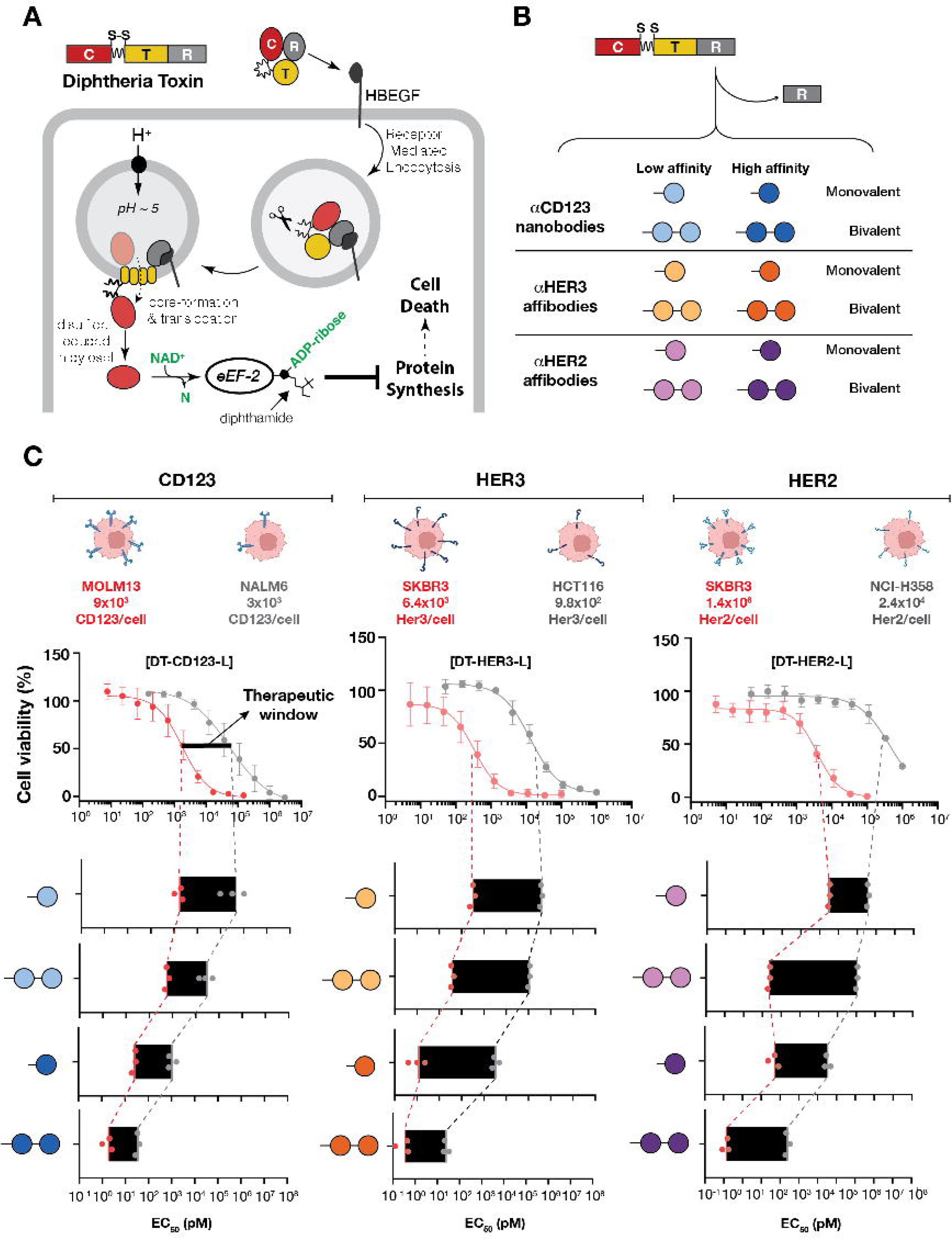
Mechanism of action of DT-based immunotoxins and the effects of affinity and avidity tuning on therapeutic window. (A). DT intoxication pathway. DT consists of a catalytic domain, a translocation domain and a receptor binding domain. Upon binding to its targeting receptor HBEGF, DT undergoes receptor-mediated endocytosis. The translocation domain facilitates endosomal escape and cytosolic delivery of the catalytic domain, which subsequently catalyzes the ADP ribosylation of elongation factor 2. This leads to inhibition of protein synthesis and ultimately cell death. (B). Schematic of DT-based immunotoxins. The native RBD of DT is swapped with a binder to cancer-associated receptors. CD123-, HER3- and HER2-targeted ITs are designed using either nanobodies (CD123) or affibodies (HER2 and HER3). For each targeted receptor, low- and high-affinity binders were formatted as either monovalent or bivalent RBDs, generating four different RBD configurations. (C). The impacts of affinity and avidity tuning on the therapeutic window of ITs. For each targeted receptor, ITs were tested on in paired receptor-high (red) and a receptor-low (gray) cell lines, with receptor abundance quantified by flow cytometry and indicated above each cell line. Representative dose-response curves are shown for the low-affinity monovalent ITs. The therapeutic window was defined as the ratio of EC50 in receptor-low over receptor-high cells. Black bars indicate the magnitude of therapeutic window, and the edge of the black bar represents the EC50 means. n=3, biological replicates.

Despite this clinical success, extending ITs to solid tumors has proven challenging, and none have received regulatory approval to date. A major limitation of ITs is their intrinsically narrow therapeutic window, driven in part by toxicity toward healthy tissues^12,13^. Most cancer-associated receptors are not strictly tumor-restricted but are also expressed, albeit at low levels in healthy cells. Consequently, even receptor-low cells can internalize small amounts of an IT and succumb to its exceptionally potent catalytic payload. One strategy to improve the therapeutic window of targeted therapies, including ITs, is to modulate the binding affinity and valency of the receptor binding domains (RBDs) to reduce binding on receptor-low cells^13,14^. However, this approach comes with an inherent trade-off: increasing discrimination against receptor-low healthy cells can also compromise activity against tumors with intermediate receptor expression.

In addition to on-target/off-tumor toxicity driven by receptor expression levels, the potency of the cytotoxic payload itself may represent an important independent determinant of the therapeutic window. This principle is well established for ADCs, where the maximum tolerated dose appears to be dictated more by payload class than by the receptor being targeted^15–17^. Payload potency may be particularly consequential for ITs because their catalytic domains are exceptionally potent. As demonstrated in a landmark study nearly 50 years ago, one molecule of DT-C introduced into a cell is sufficient to kill that cell^18^. Thus, even limited uptake by receptor-low/-null healthy cells may deliver sufficient toxin to exceed the threshold required for cytotoxicity. We therefore reasoned that tuning receptor binding alone may be insufficient to optimize the therapeutic window of ITs, and that attenuating the catalytic activity of the payload could preferentially spare receptor-low cells while preserving activity against receptor-high tumor cells.

In this work, we asked whether the therapeutic window of immunotoxins could be expanded by tuning not only receptor engagement, but also the catalytic potency of the toxin payload. Using CD123-, HER3-, and HER2-targeted immunotoxins, we systematically varied receptor-binding affinity, avidity, and catalytic activity to define how each contributes to discrimination between receptor-high and receptor-low cells. Unexpectedly, this analysis revealed catalytic potency as a major and independently tunable determinant of immunotoxin selectivity. More broadly, our findings uncover a design principle in which optimal immunotoxin activity is achieved not by maximizing receptor binding or payload potency, but by appropriately matching the two.

## Results

### Affinity and avidity tune immunotoxin potency but provide limited control over selectivity

To determine how receptor-binding affinity and avidity influence immunotoxin potency and selectivity, we generated a systematic panel of diphtheria toxin (DT)-based ITs targeting three cancer-associated receptors with distinct expression profiles: CD123, HER3, and HER2. Each IT comprised the catalytic and translocation domains of DT (DT_1–389_), with the native receptor-binding domain replaced by a target-specific nanobody or affibody (**Fig. 1A and B**). For each receptor, we selected previously characterized low- and high-affinity binders and formatted each as either a monovalent or bivalent receptor-binding domain, generating four binding configurations that independently varied affinity and avidity (**Fig. 1B**). We next quantified receptor abundance on the corresponding receptor-high and receptor-low cell lines by flow cytometry. These measurements revealed distinct differences in receptor abundance between receptor-high and receptor-low cell lines across the three targets, with HER2 showing a substantially greater separation than either CD123 or HER3.

We evaluated each IT series in pairs of cell lines differing in target-receptor abundance. CD123-targeted ITs were tested in MOLM13 and NALM6 cells, which expressed approximately 9 × 10^3^ and 3 ×10^3^ CD123 molecules per cell, respectively. HER3-targeted ITs were evaluated in SK-BR-3 and HCT116 cells, expressing 6.4 × 10^3^ and 9.8 × 10^2^ HER3 molecules per cell, while HER2-targeted ITs were tested in SK-BR-3 and NCI-H358 cells, expressing 1.4 × 10^6^ and 2.4 × 10^4^ HER2 molecules per cell, respectively (**Fig. 1C**). We define the **therapeutic window** here as the ratio of EC_50_ values between the receptor-low and receptor-high cell lines.

Across all three receptors, increasing RBD affinity and/or avidity substantially increased IT potency. However, these potency shifts were generally observed in both receptor-high and receptor-low cells, such that improvements in receptor engagement did not consistently translate into proportional gains in selectivity (**Fig. 1C**). For example, the low-affinity monovalent CD123-targeted IT was more potent in CD123-high MOLM13 cells than in CD123-low NALM6 cells, but increasing affinity or avidity shifted potency toward lower concentrations in both cell types. Similar behavior was observed across the HER3- and HER2-targeted series despite marked differences in absolute receptor densities and differential receptor expression across these systems.

Thus, although affinity and avidity are effective determinants of IT potency, modulating receptor engagement alone provides limited control over the separation between receptor-high and receptor-low cells. The parallel shifts in potency across receptor-high and receptor-low cells raised the possibility that the extreme activity of the WT catalytic payload limits how effectively differences in receptor-mediated uptake can be translated into selective cytotoxicity.

### Catalytic activity tuning dramatically expands immunotoxin selectivity

DT-C is an ADP-ribosyltransferase that inhibits protein synthesis through covalent modification of eEF-2^19,20^. Its catalytic activity is exceptionally high, with as little as a single molecule of DT-C in the cytosol reported to be sufficient to kill a cell^18^. We therefore hypothesized that the high intrinsic activity of WT DT-C may limit the selectivity achievable through receptor-binding optimization. Specifically, we reasoned that cell death requires inactivation of a critical threshold of eEF-2 and, consequently, delivery of a sufficient amount of active DT-C. Reducing catalytic activity should increase the amount of intracellular payload required to reach this lethal threshold. Under this model, greater uptake in receptor-high cells could remain sufficient for killing, whereas the lower levels of uptake in receptor-low cells could fall below the threshold required for cytotoxicity.

To test this hypothesis, we introduced two previously characterized active-site mutations into the DT catalytic domain: K51E, which retains approximately 10% of WT activity, and E148D, which retains approximately 1.6% of WT activity^21^. E148 is a key residue involved in catalyzing ADP-ribosyl transfer to EF-2^22^. K51 is involved in NAD^+^ binding^23^. These mutations were introduced across the low- and high-affinity, monovalent and bivalent RBD formats described above, allowing us to determine how catalytic activity interacts with receptor engagement to control selectivity.

To determine how catalytic activity interacts with receptor engagement, we first compared WT, K51E, and E148D payloads across the four HER2-targeting RBD formats (**Fig. 2**). In the low-affinity RBD formats, reducing catalytic activity produced little improvement in the therapeutic window, as attenuation reduced potency in HER2-high and HER2-low cells to similar extents. In contrast, attenuation substantially increased selectivity in the high-affinity formats. Most notably, reducing catalytic activity to 1.6% in the high-affinity bivalent construct, DT^E148D^-HER2_H_×2, preserved potent cytotoxicity toward HER2^+++^ SK-BR-3 cells while eliminating detectable toxicity toward HER2^+^ NCI-H358 cells over the concentration range tested. This increased the therapeutic window to approximately 4.6 × 10^6^, representing nearly a 2,500-fold improvement relative to the corresponding WT IT. Because no EC50 could be determined in NCI-H358 cells, this therapeutic window represents a lower-bound estimate. Importantly, the relationship between catalytic activity and selectivity was not monotonic: the intermediate K51E mutation did not improve selectivity in every RBD format, indicating that attenuation alone is insufficient and that payload activity must be appropriately matched to receptor engagement.

**Figure 2.**
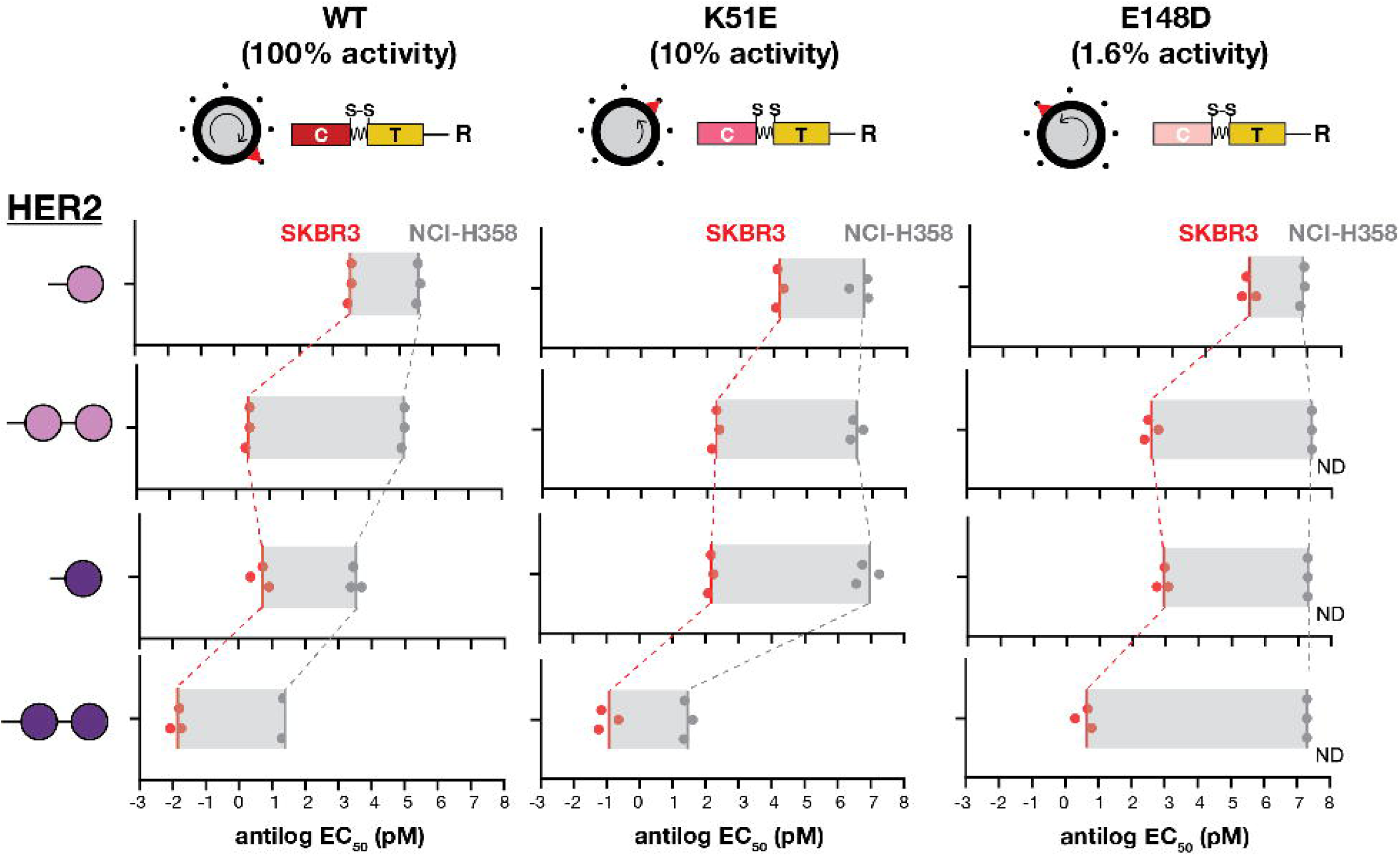
The combine effects of affinity, avidity and activity tuning of HER2-targeted ITs. HER2-targeted ITs with WT DT-C (100% activity), K51E (around 10% activity) or E148D (around 1.6% activity) were evaluated across four HER2 RBD formats. Cytotoxicity was measured on HER2^+++^ SK-BR-3 (red) and HER2^+^ NCI-H358 (gray) cells. Each point represents EC50 values from independent experiments, and the edge of the gray bar indicates the EC50 means (n=3, biological replicates). Gray bars indicate the magnitude of therapeutic window. The EC50s of WT ITs were also presented on Figure 1. ND: EC50 not determined because 50% cytotoxicity was not reached.

Similar behavior was observed with CD123- and HER3-targeted ITs (**Supplementary Fig. 1**). For both receptors, catalytic attenuation produced the largest gains in selectivity in the high-affinity bivalent formats. However, in contrast to HER2, maximal selectivity was achieved with the intermediate K51E mutation rather than E148D. For CD123, K51E increased the selectivity of the high-affinity bivalent IT more than 5,000-fold, whereas for HER3 the corresponding increase exceeded 30,000-fold. Further attenuation with E148D reduced on-target potency and consequently decreased selectivity. Consistent with the proposed threshold model, the increased selectivity of tuned ITs was associated with loss of detectable receptor-dependent cytotoxicity in receptor-low cells. For example, DT^K51E^-CD123_H_×2 was equipotent to an untargeted DT^K51E^ control in NALM6 cells (**Supplementary Fig. 2**), indicating that low-level CD123-mediated uptake was no longer sufficient to support detectable cytotoxicity. By contrast, the corresponding WT IT retained substantial receptor-dependent activity in these cells.

Together, these results show that reducing catalytic activity does not uniformly improve selectivity. Rather, maximal selectivity emerges at an intermediate payload activity that preserves killing when receptor-mediated uptake is high but is insufficient to reach lethal threshold when uptake is limited. The optimal catalytic activity therefore depends on the strength of receptor engagement.

### Payload activity determines whether differential receptor engagement translates into selective cytotoxicity

The preceding experiments showed that maximal selectivity depends on matching catalytic activity to receptor engagement. We next asked whether activity tuning could generate selective cytotoxicity when differences in receptor abundance were relatively small. To address this, we compared HER2^++^ JIMT-1 cells, which express approximately 4.8 × 10^4^ HER2 molecules per cell, with HER2^+^ NCI-H358 cells, which express approximately 2.4 × 10^4^ receptors per cell, an approximately two-fold difference (**Fig. 3A**).

**Figure 3.**
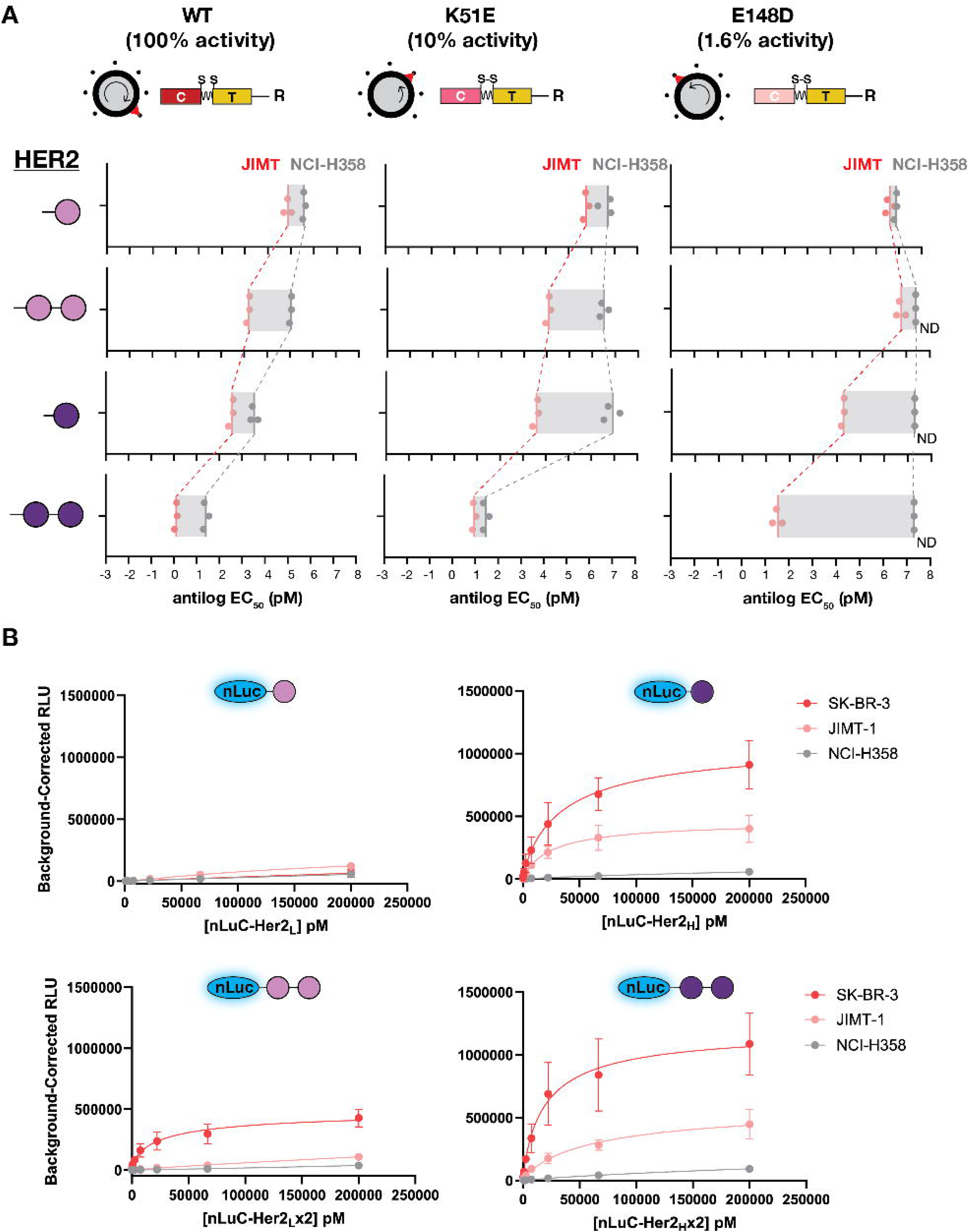
The effects of catalytic activity tuning on HER2-targeted ITs on moderate HER2 expressing cells. (A). The therapeutic window of HER2-targeted ITs on HER2^++^ cells. HER2-targeted ITs with WT DT-C (100% activity), K51E (around 10% activity) or E148D (around 1.6% activity) were evaluated across four HER2 RBD formats. Cytotoxicity was measured on HER2^++^ JIMT-1 (pink) and HER2^+^ NCI-H358 (gray) cells. Each point represents EC50 values from independent experiments, and the edge of the gray bar indicates the EC50 means (n=3, biological replicates). Gray bars indicate the magnitude of therapeutic window. The EC50s of ITs on NCI-H358 were also presented on Figure 2. ND: EC50 not determined because 50% cytotoxicity was not reached. (B). On cell binding of HER2 RBDs across three cell lines with distinct HER2 expression. HER2 RBDs were fused to NanoLuc (nLuc), and cell binding was quantified as background-corrected luminance. No cell control was used as the background signal. n=3, biological replicates.

Across the four HER2-targeting RBD formats, the effects of catalytic attenuation followed a similar pattern to those observed with HER2^+++^ SK-BR-3 cells. In the low-affinity formats, reducing payload activity provided little improvement in selectivity, as attenuation also compromised cytotoxicity toward JIMT-1 cells. In contrast, activity tuning substantially increased selectivity in the high-affinity formats. Most notably, DT^E148D^-HER2_H_×2 retained potent activity against JIMT-1 cells while showing no detectable toxicity toward NCI-H358 cells over the concentration range tested, resulting in a therapeutic window of approximately 5.8 × 10^5^ and an almost 30,000-fold improvement over its WT counterpart (**Fig. 3A**). Thus, appropriately attenuated payloads could translate even relatively modest differences in receptor abundance into large differences in cytotoxic response.

We next asked whether this selectivity could be explained simply by preferential receptor binding. To directly measure cell-surface engagement, we fused each HER2 RBD to NanoLuc and quantified binding to SK-BR-3, JIMT-1, and NCI-H358 cells (**Fig. 3B**). Increasing affinity and avidity enhanced cell-surface binding and, particularly for the bivalent formats, produced preferential engagement of HER2-high cells relative to NCI-H358 cells. However, these differences in receptor engagement were not sufficient to generate comparable differences in cytotoxicity when the RBDs were paired with WT DT-C. In particular, the high-affinity formats showed clear preferential binding to HER2-higher cells yet remained only modestly selective when coupled to the WT payload.

These findings indicate that differential receptor engagement alone does not determine cell-type selectivity. Rather, payload activity determines whether differences in receptor engagement are translated into differences in cell fate. With the highly active WT payload, even limited receptor-mediated uptake by receptor-low cells appears sufficient to support cytotoxicity. Catalytic attenuation raises the amount of intracellular toxin required for killing, allowing greater receptor engagement in cells with higher HER2 expression cells to remain productive while lower levels of uptake in receptor-low cells fall below the lethal threshold. Thus, maximal selectivity requires payload activity to be matched to the magnitude of receptor engagement.

### Catalytic activity tuning extends across toxin scaffolds

Having established that catalytic attenuation can markedly improve the selectivity of DT-based immunotoxins, we next asked whether this principle extends to other toxin scaffolds. To test this, we used Chelona toxin (CT), a diphtheria toxin-like toxin from *Austwickia chelonae*^24^. We generated a HER2-targeted CT immunotoxin containing the CT2 catalytic domain and CT1 translocation domain fused to the high-affinity bivalent HER2 RBD, HER2_H_×2, which produced the greatest selectivity in the DT series.

To generate a comparable range of catalytic activities, we introduced mutations in CT corresponding to those used in DT-C: K51E, E148D, and W50R. We then compared the selectivity of these constructs across HER2^+++^ SK-BR-3, HER2^++^ JIMT-1, and HER2^+^ NCI-H358 cells (**Fig. 4**). Attenuating CT catalytic activity also markedly increased immunotoxin selectivity. The K51E variant produced the greatest improvement, increasing selectivity by more than 10,000-fold relative to the WT CT payload. In contrast, further attenuation with E148D reduced selectivity, while W50R abolished detectable activity in HER2^+++^ and HER2^++^ cells. Notably, the optimal level of catalytic attenuation differed from that observed with DT, where E148D produced the greatest selectivity in the corresponding HER2_H_×2 format.

**Figure 4.**
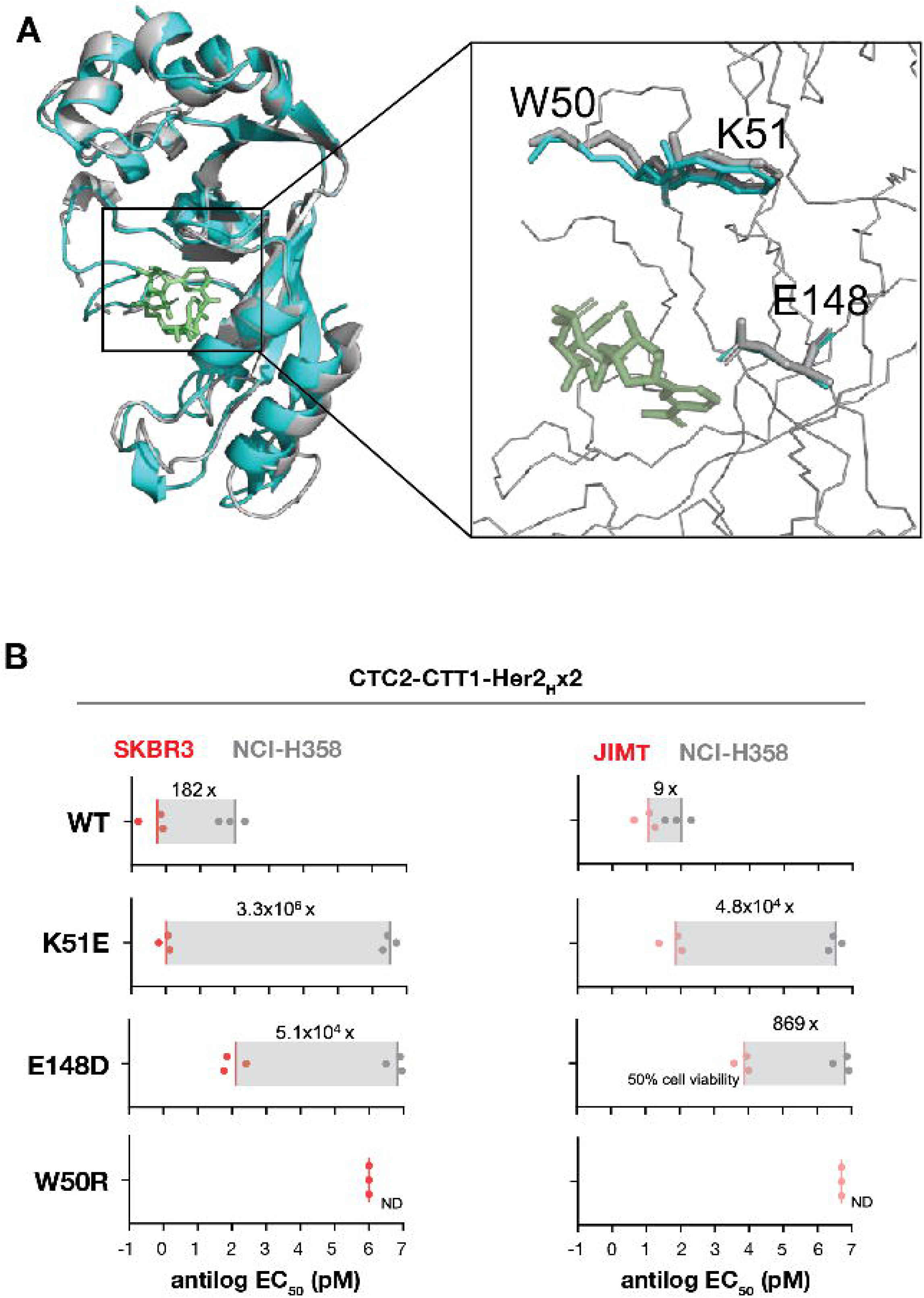
Activity tuning expands the therapeutic window of CT-based HER2-targeted ITs. (A). Structural comparison of the DT-C and CT-C2. The catalytic domains of DT (PDB 1TOX) and CT2 were structurally aligned, with the catalytic residues W50, K51 and E148D highlighted. NAD is shown in green. CT-C structure was generated using AlphaFold3 webserver. (B). The effects of activity tuning on CT-based HER2-targeted ITs. CTC2-CTT1-HER2_H_x2 was used as a template. WT CT-C, K51E, E148D or W50R were evaluated on HER2^+++^ SK-BR-3 versus HER2^+^ NCI-H358 (left) and HER2^++^ JIMT-1 versus HER2^+^ NCI-H358 cells (right). Each point represents EC50 values from independent experiments, and the edge of the gray bar indicates the EC50 means (n=3, biological replicates). Gray bars indicate the magnitude of therapeutic window. ND: EC50 not determined because 50% cytotoxicity was not reached.

These results demonstrate that catalytic activity is a tunable determinant of selectivity across distinct toxin scaffolds. Importantly, however, the optimal payload activity is not defined by a particular mutation or fixed level of attenuation, but depends on the properties of the toxin scaffold and its interaction with receptor-mediated delivery. Together, these findings support a broader design principle in which immunotoxin selectivity is maximized by matching catalytic payload activity to the efficiency of cellular delivery.

### Payload attenuation creates an *in vivo* therapeutic window for antitumor efficacy

To determine whether the large gains in *in vitro* selectivity achieved by catalytic attenuation translate into an improved therapeutic window *in vivo*, we compared the HER2-targeted WT and E148D ITs in mice. We selected the high-affinity bivalent HER2_H_×2 format because E148D produced the greatest increase in selectivity in this context. An albumin-binding domain (ABD) was fused to both constructs to extend their circulating half-life. For efficacy studies, we used NCI-N87 xenografts, which express high levels of HER2 and reliably form tumors in mice.

We first determined the maximum tolerated dose (MTD) of each construct in healthy mice. The attenuated DT^E148D^-HER2_H_×2 IT was tolerated at doses up to 20 mg/kg, whereas the corresponding WT IT was tolerated only up to 0.05 mg/kg, representing an approximately 400-fold increase in tolerated dose following catalytic attenuation (**Supplementary Fig. 3**). Thus, the substantial increase in cellular selectivity produced by activity tuning was accompanied by a similarly pronounced improvement in *in vivo* tolerability.

We next asked whether this expanded tolerability translated into a therapeutically useful dose range. Because the WT and E148D constructs differed markedly in their MTDs, tumor-bearing mice were treated at matched fractions of their respective MTDs: 1/64, 1/16, and 1/4, allowing efficacy to be compared relative to each molecule’s toxicity ceiling (**Fig. 5**). At 1/64 of the MTD, neither construct substantially inhibited tumor growth. In contrast, DT^E148D^-HER2_H_×2 produced progressively greater tumor growth inhibition at 1/16 and 1/4 of its MTD, whereas the WT IT showed little to no antitumor activity even at 1/4 of its MTD.

**Figure 5.**
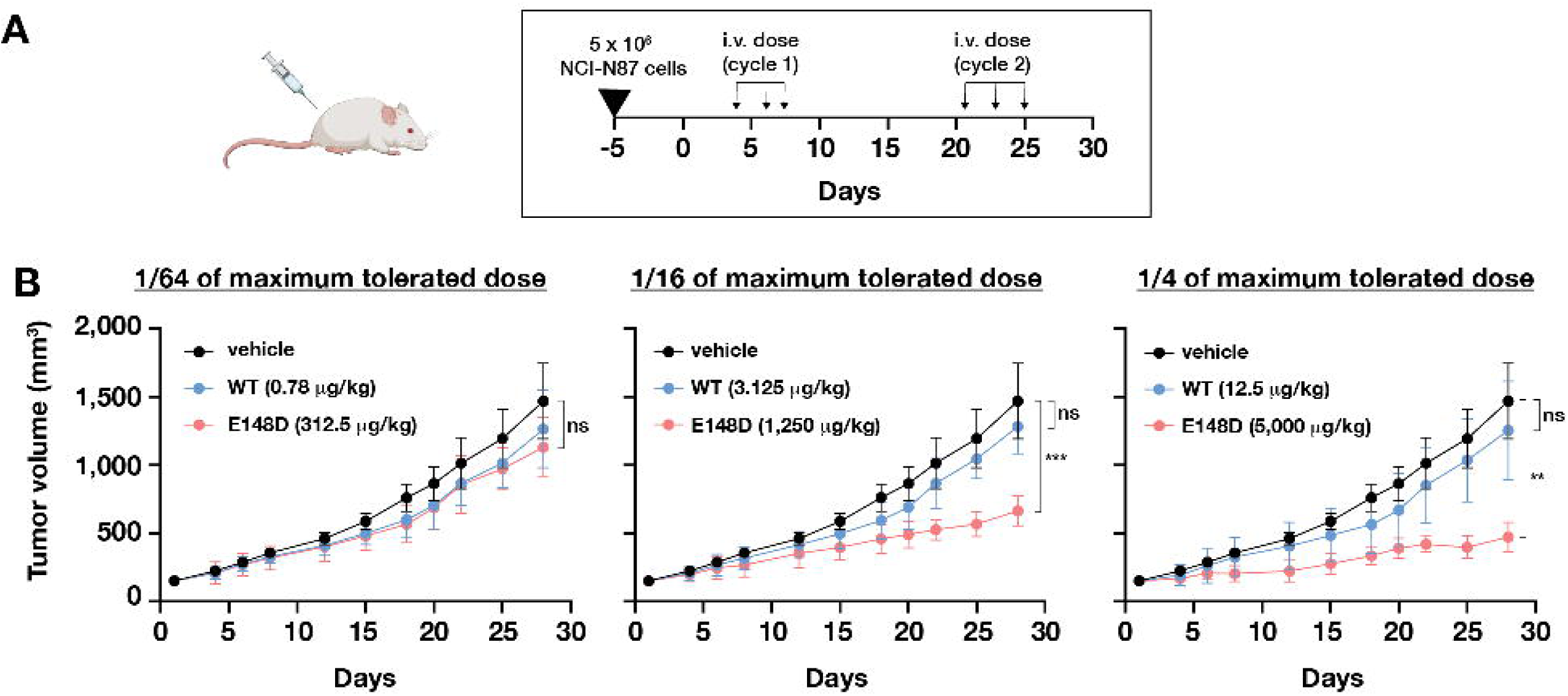
Activity tuning enables anti-tumor activity *in vivo*. (A). Schematic of the NCI-N87 xenograft efficacy study. Female NOD SCID mice were subcutaneously inoculated with 5 × 10⁶ NCI-N87 cells with treated intravenously with vehicle, DT-HER2_H_x2-ABD or DT^E148D^-HER2_H_x2-ABD on indicated days. (B). Anti-tumor efficacy of HER2-targeted ITs. Tumor-bearing mice were treated at matched fractions of their respective maximal tolerated doses (1/64, 1/16 and 1/4). Tumor volumes were monitored throughout the study. n=7. ns, not significant; **P < 0.01; ***P < 0.001.

These results demonstrate that catalytic attenuation does more than simply increase the amount of IT that can be administered. By widening the separation between efficacious and toxic exposures, attenuation creates a therapeutically accessible dose range that is largely absent for the WT payload. Thus, reducing catalytic activity can convert an IT whose systemic toxicity prevents effective dosing into one capable of achieving antitumor activity well below its toxicity limit.

## Discussion

Immunotoxins (ITs) combine selective receptor targeting with exceptionally potent catalytic payloads, but their broader clinical application has been constrained by narrow therapeutic windows and dose-limiting toxicity^13^. A central challenge for all targeted therapies, including ITs, is that tumor-associated receptors are rarely restricted to malignant cells. Thus, even low levels of receptor expression on normal tissues can permit sufficient IT uptake to cause toxicity. Our findings uncovered catalytic payload activity as a major determinant of this selectivity. Across CD123-, HER3-, and HER2-targeted ITs, altering RBD affinity and avidity changed potency but provided limited and inconsistent gains in discrimination between receptor-high and receptor-low cells. In contrast, appropriately attenuating catalytic activity produced increases in selectivity of several orders of magnitude. These results support a model in which IT selectivity is determined not by receptor engagement or payload potency alone, but by the relationship between cellular delivery and the catalytic activity required to trigger cell death.

Previous approaches to improving the selectivity of targeted therapeutics have largely focused on receptor engagement^13,14^. Affinity tuning can exploit differences in receptor density by favoring binding to receptor-high tumor cells while reducing engagement of receptor-low normal cells. For example, Jun and colleagues showed that intermediate-affinity EGFR-targeted ITs provided greater selectivity than either high-or low-affinity variants^14^. Related principles have also been demonstrated with affinity-tuned CAR-T cells targeting EGFR and HER2^25^. Our results are consistent with receptor affinity and avidity being important determinants of IT potency, but show that differential receptor engagement does not necessarily translate into comparable differences in cytotoxicity. Across three receptors with markedly different expression profiles, changing RBD affinity and avidity frequently shifted potency in receptor-high and receptor-low cells in parallel. Moreover, HER2-targeted ITs with substantially different *in vitro* selectivity nevertheless exhibited similarly poor tolerability when paired with WT DT-C. Together, these findings indicate that optimization of receptor engagement alone may be insufficient when the downstream payload is exceptionally potent.

The extraordinary catalytic activity of DT provides a mechanistic explanation for this limitation. A landmark study demonstrated that delivery of very few, and potentially a single, DT catalytic-domain molecules to the cytosol can be sufficient to kill a cell^18^. Under these conditions, relatively small differences in receptor-mediated delivery may have little consequence for cell fate: both receptor-high and receptor-low cells may receive enough active toxin to exceed the lethal threshold. Attenuating catalytic activity changes this relationship by increasing the amount of intracellular payload required to kill a cell. Differences in receptor engagement can then become biologically consequential, because uptake by receptor-low cells may fall below the threshold for cytotoxicity while greater uptake by receptor-high cells remains sufficient for killing. Consistent with this model, differential HER2 binding measured directly on cells did not generate comparable differences in cytotoxicity when the RBDs were coupled to WT DT-C, whereas catalytic attenuation converted these differences in receptor engagement into pronounced differences in cell killing.

Importantly, reducing payload activity did not uniformly improve selectivity. The largest gains occurred when attenuated catalytic domains were paired with RBD formats that provided sufficiently strong receptor engagement. Excessive attenuation compromised killing of receptor-high cells, whereas insufficient attenuation retained toxicity toward receptor-low cells. Thus, maximal selectivity occurred within an activity window in which receptor-mediated delivery resulted in sufficient amount of toxin to kill target cells but fell below the lethal threshold in receptor-low cells. The position of this window varied among CD123, HER3, and HER2 and also between DT and CT. For example, the E148D mutation provided optimal selectivity for HER2-targeted DT constructs, whereas the corresponding degree of attenuation was suboptimal in the CT scaffold. These findings indicate that there is no universally optimal catalytic activity; instead, payload potency must be matched empirically to receptor abundance, RBD architecture, and the intracellular delivery efficiency of the toxin scaffold.

The CT experiments further suggest that activity tuning represents a property of immunotoxin design rather than a peculiarity of DT. This distinction is important because it argues against individual mutations themselves being responsible for the effect. Rather, the relevant variable is whether the amount and activity of toxin delivered to the cytosol are sufficient to exceed the threshold required for cell killing. In this framework, receptor binding, internalization, translocation, and catalytic activity collectively determine whether a given cell crosses the cytotoxic threshold.

The therapeutic consequence of this principle was evident *in vivo*. The activity-tuned HER2-targeted IT, DT^E148D^-HER2_H_×2, exhibited an approximately 400-fold higher maximum tolerated dose than its WT counterpart. More importantly, this improved tolerability translated into a therapeutically accessible dose range. DT^E148D^-HER2_H_×2 produced robust tumor growth inhibition at fractions of its MTD, including activity at one-sixteenth of the MTD, whereas the WT construct showed little detectable efficacy at the tolerated doses tested. Thus, catalytic attenuation did not simply permit administration of more drug; it increased the separation between efficacious and toxic exposures. This is the central pharmacological consequence of activity tuning: a highly potent payload can generate substantial cytotoxic activity but leave little room between tumor killing and systemic toxicity, whereas appropriately attenuating that payload can create a usable therapeutic window.

These findings suggest a broader model for the design of targeted cytotoxic agents. The optimal payload should not necessarily be the most active payload available. Instead, its activity should be matched to the magnitude of differential delivery between target and non-target cells. When receptor expression differs greatly between tumor and normal tissue, greater attenuation may be tolerated while preserving tumor-cell killing. When that difference is smaller, the acceptable activity window may narrow. Receptor abundance alone, however, is unlikely to fully predict this optimum because cellular delivery also depends on receptor internalization, intracellular trafficking, translocation efficiency, and payload stability. These parameters will ultimately determine the amount of active payload reaching its intracellular target.

Several limitations should therefore be considered. We quantified receptor abundance and cell-surface engagement but did not independently measure internalization, cytosolic delivery, or the number of active toxin molecules reaching eEF-2. The proposed threshold model is therefore strongly supported by the pharmacological behavior of the constructs but remains to be tested directly. In addition, the *in vivo* efficacy studies were performed in a single HER2-high xenograft model. Evaluation across additional tumor models, particularly those with more moderate HER2 expression, together with detailed pharmacokinetic and tissue-distribution studies, will be important for defining how catalytic attenuation influences exposure, efficacy, and safety *in vivo*. Nevertheless, the consistent effects observed across three receptors and two toxin scaffolds, together with the marked improvement in tolerability and antitumor activity, support payload tuning as a general strategy for widening the therapeutic window of ITs.

We contend that this principle could extend beyond immunotoxins. Other targeted therapeutics couple selective cellular recognition to highly active downstream effectors, and excessive effector activity may similarly obscure quantitative differences in target engagement. For example, reduced protease activity was recently required to achieve selective degradation of a tumor-associated surface glycan while sparing normal cells^26^. More broadly, our results argue that maximal molecular activity is not necessarily synonymous with maximal therapeutic performance. Matching payload activity to cellular delivery may provide a general strategy for converting quantitative differences between target and normal cells into a wider therapeutic window.

## Methods

### Cell cultures

Human breast adenocarcinoma cell line SK-BR-3, human colorectal cancer cell line HCT116, human non-small cell lung cancer cell line NCI-H358, human acute myeloid leukemia cell line MOLM13, and human acute lymphoblastic leukemia cell line NALM6 were purchased from ATCC. Human breast carcinoma cell line JIMT-1 was purchased from AddexBio. SK-BR-3, NCI-H358, MOLM13 and NALM6 were cultured in RPMI1640 media. JIMT-1 was cultured in DMEM media. HCT116 was cultured in McCoy’s media. All media was supplemented with 10% heat-inactivated FBS and 1% penicillin-streptomycin. All cell lines were maintained at 37 °C with 5% CO_2_.

### Plasmid design and construction

The gene fragments of the receptor binding domains were synthesized as gblocks. All plasmids contain a N-terminal His_6_-SUMO tag, and a C-terminal Strep-tag II.

DT-based immunotoxins were generated through genetically fusing various receptor binding domains to DT1-389 in a pETSUMO vector. A (G_4_S)_2_ linker was inserted between DT fragment and the receptor binding domain. CT-based immunotoxins were constructed through genetically fused CTC2-CTT1 with receptor binding domains. For bivalent receptor binding domain design, there is a (G_4_S)_3_ linker between the individual receptor binding domain. Vhh55A01 and Vhh57A07 were used as a receptor binding domain to CD123. ZHER2:4, ZHER2:342 were used as receptor binding domains to HER2. ZHER3:01820 and ZHER3:08699 were used as a receptor binding domain to HER3. ZTaq was used as a dummy receptor binding domain for the untargeted control. Individual point mutations in the DT-C domain were introduced through QuikChange Lightning Multi-site directed mutagenesis, following the manufactural protocol. ABD fusion ITs were constructed by genetically fusing ABD035 to the C terminus of DT-HER2_H_x2 and DT^E148D^-HER2_H_x2 with a (G_4_S)_3_ linker.

### Protein expression and purification

Plasmids were transformed into *E.coli* Nico21 cells (NEB). Single clone was picked and grown overnight in 50mL of LB broth with 50µg/mL kanamycin at 37 °C. The next day, overnight was inoculated into 1L of LB containing 50µg/mL kanamycin at 37 °C. Protein expression was induced when OD600 reached ∼0.7 with 1mM IPTG, and cells were harvested after 5 hours of induction at 25 °C through centrifugation at 5,000g for 20 min. Cell pellets were resuspended in 30mL of lysis buffer (20mM Tris-HCl pH 8, 300mM NaCl, 1% protease inhibitor cocktail, 0.01% universal nuclease) and lysed by sonication. The whole cell lysates were centrifuged at 18,000g for 20 min to isolate the soluble fraction. The soluble fraction was loaded onto a 5mL Ni-Penta Agarose 6FF column using AKTA GO purification system. The recombinant proteins with His6 tag were eluted with elution buffer (20mM Tris-HCl pH 8, 300mM NaCl, 500mM imidazole), and the elution fractions were subject to SUMO protease treatment overnight at 4 °C. The proteins were then loaded onto a 5mL Strep-Tactin XT column using AKTA GO purification system, and eluted with elution buffer (20mM Tris-HCl pH 8, 150mM NaCl, 50mM biotin). The elution was further concentrated using spin concentrators (Millipore). Glycerol was added to the concentrated solution to achieve 5% glycerol. Finally, the proteins were sterilized with 0.22µm centrifuge tube filters (Costar). Protein purity was assessed by SDS-PAGE gels and the concentration was measured using a NanoDrop. For *in vivo* usage, endotoxin was further removed from the protein solutions using EndoTrap HP column (Lionex) and endotoxin level was quantified by LAL quantification kit (Charles River Laboratories).

### Cell viability

Cells were trypsinized and seeded onto a 96-well plate (Corning) at 4-7,000 cells/well. The next day, ITs were prepared as three-fold serial dilutions at 10x final concentration and added to cells to achieve the indicated final concentrations. Cells were incubated with immunotoxins for 72 hours and cell viability was measured using PrestoBlue reagent following the manufactory protocol. The fluorescence reading was measured using SpectraMax M5e microplate reader (excitation at 555nm, filter at 575nm, emission at 585nm). The data was analyzed using prism.

### Receptor quantification

CD123, HER2 and HER3 receptor densities were quantified by Quantum R-PE MESF (Bang’s Laboratory) following the manufactory protocol. Briefly, cells were seeded on 12-well plates at 80% confluency. The next day, cells were stained with PE-conjugated antibodies on ice for 1 hour at saturating concentration for each individual cell line. Cells were washed with cold wash buffer (PBS, 2% BSA) and detached from the plates using non-enzymatic cell dissociation solution. Non-bound antibody was removed through centrifuge and washed 3x with wash buffer. For suspension cells, no detachment was performed, and excessive antibody was removed through centrifugation and washed for 3 times. Cells were then analyzed by flow cytometry along with Bang Laboratories’ Quantum R-PE MESF standard beads. MFI was analyzed using FlowJo, and receptor density was calculated based on the MFI values generated from the standard curve. Anti-CD123, anti-HER2 and anti-HER3 antibody were used for cell staining.

### Maximal tolerated dose study

NOD SCID mice (n=3) were administrated intravenously with DT^E148D^-HER2_H_x2-ABD or DT-HER2_H_x2-ABD for total of three doses. Body weight was monitored daily. A dose was considered non-tolerated if it resulted in greater than 20% weight loss that did not recover after dosing was discontinued. The MTD was defined as the highest dose that did not meet this criterion. The study was performed by Pharmaron.

### NCI-N87 xenograft experiment

Female NODSCID mice (6-8 weeks old) were subcutaneously inoculated in the right flank with 5×10^6^ NCI-N87 cells suspended in 100µL of a medium/Matrigel mixture. Treatment was initiated when the tumor volume was greater than 150mm^3^. ITs were diluted in vehicle buffer (20mM Tris pH 8, 150mM NaCl), and were administered intravenously for a total of 6 doses (3 doses/cycle, totally 2 cycles). Tumor volume and body weight were monitored during the study.

## Acknowledgments and funding sources

The authors thank members of the Melnyk lab for critical comments on this manuscript. This work was supported by grants from the Canadian Institutes of Health Research (CIHR) (to R.A.M.) (#452521), and the Natural Sciences and Engineering Research Council of Canada RGPIN-2023-05371 (to R.A.M.).

## Declaration of Interests

R.A.M. is a scientific advisory board member for Adaxion Therapeutics. R.A.M, H.L, and G.LB. have patent applications (issued and pending) related to this work

## Notes

### Competing Interest Statement

HL, GLB and RAM have filed provisional patents related to this work.

## References

1. Zhu, K. et al. HER2-targeted therapies in cancer: a systematic review. Biomarker Research 12, 16 (2024).

2. Tebbutt, N., Pedersen, M. W. & Johns, T. G. Targeting the ERBB family in cancer: couples therapy. Nat Rev Cancer 13, 663–673 (2013).

3. Gandullo-Sánchez, L., Ocaña, A. & Pandiella, A. HER3 in cancer: from the bench to the bedside. J Exp Clin Cancer Res 41, 310 (2022).

4. Seshacharyulu, P. et al. Targeting the EGFR signaling pathway in cancer therapy. Expert Opin Ther Targets 16, 15–31 (2012).

5. Oh, D.-Y. & Bang, Y.-J. HER2-targeted therapies — a role beyond breast cancer. Nat Rev Clin Oncol 17, 33–48 (2020).

6. Allen, T. M. Ligand-targeted therapeutics in anticancer therapy. Nat Rev Cancer 2, 750–763 (2002).

7. Weldon, J. E. & Pastan, I. A guide to taming a toxin: recombinant immunotoxins constructed from Pseudomonas exotoxin A for the treatment of cancer. FEBS J 278, 4683– 4700 (2011).

8. Shafiee, F., Aucoin, M. G. & Jahanian-Najafabadi, A. Targeted Diphtheria Toxin-Based Therapy: A Review Article. Front Microbiol 10, 2340 (2019).

9. Duvic, M. & Talpur, R. Optimizing Denileukin Diftitox (Ontak®) Therapy. Future Oncology 4, 457–469 (2008).

10. Pemmaraju, N. & Konopleva, M. Approval of tagraxofusp-erzs for blastic plasmacytoid dendritic cell neoplasm. Blood Advances 4, 4020–4027 (2020).

11. Dhillon, S. Moxetumomab Pasudotox: First Global Approval. Drugs 78, 1763–1767 (2018).

12. Alewine, C., Hassan, R. & Pastan, I. Advances in Anticancer Immunotoxin Therapy. Oncologist 20, 176–185 (2015).

13. Li, M., Mei, S., Yang, Y., Shen, Y. & Chen, L. Strategies to mitigate the on- and off-target toxicities of recombinant immunotoxins: an antibody engineering perspective. Antibody Therapeutics 5, 164–176 (2022).

14. Jun, S.-Y., Kim, D.-S. & Kim, Y.-S. Expanding the Therapeutic Window of EGFR-Targeted PE24 Immunotoxin for EGFR-Overexpressing Cancers by Tailoring the EGFR Binding Affinity. Int J Mol Sci 23, 15820 (2022).

15. Colombo, R. & Rich, J. R. The therapeutic window of antibody drug conjugates: A dogma in need of revision. Cancer Cell 40, 1255–1263 (2022).

16. Rubahamya, B., Dong, S. & Thurber, G. M. Clinical translation of antibody drug conjugate dosing in solid tumors from preclinical mouse data. Sci Adv 10, eadk1894.

17. Nguyen, T. D., Bordeau, B. M. & Balthasar, J. P. Mechanisms of ADC Toxicity and Strategies to Increase ADC Tolerability. Cancers 15, 713 (2023).

18. Yamaizumi, M., Mekada, E., Uchida, T. & Okada, Y. One molecule of diphtheria toxin fragment a introduced into a cell can kill the cell. Cell 15, 245–250 (1978).

19. Collier, R. J. Understanding the mode of action of diphtheria toxin: a perspective on progress during the 20th century. Toxicon 39, 1793–1803 (2001).

20. Collier, R. J. Diphtheria toxin: mode of action and structure. Bacteriol Rev 39, 54–85 (1975).

21. Fu, H., Blanke, S. R., Mattheakis, L. C. & Collier, R. J. Selection of Diphtheria Toxin Active-Site Mutants in Yeast. in ADP-Ribosylation in Animal Tissues: Structure, Function, and Biology of Mono (ADP-ribosyl) Transferases and Related Enzymes (eds. Haag, F. & Koch-Nolte, F.) 45–52 (Springer US, Boston, MA, 1997). doi:10.1007/978-1-4419-8632-0_5.

22. Wilson, B. A., Reich, K. A., Weinstein, B. R. & Collier, R. J. Active-site mutations of diphtheria toxin: effects of replacing glutamic acid-148 with aspartic acid, glutamine, or serine. Biochemistry 29, 8643–8651 (1990).

23. Uthailak, N. et al. Molecular Cloning, Structural Modeling and the Production of Soluble Triple-Mutated Diphtheria Toxoid (K51E/G52E/E148K) Co-expressed with Molecular Chaperones in Recombinant Escherichia coli. Mol Biotechnol 59, 117–127 (2017).

24. Gill, S. K. et al. An enhanced intracellular delivery platform based on a distant diphtheria toxin homolog that evades pre-existing antitoxin antibodies. EMBO Mol Med 16, 2638– 2651 (2024).

25. Liu, X. et al. Affinity-tuned ErbB2 or EGFR chimeric antigen receptor T cells exhibit an increased therapeutic index against tumors in mice. Cancer Res 75, 3596–3607 (2015).

26. Pedram, K. et al. Design of a mucin-selective protease for targeted degradation of cancer-associated mucins. Nat Biotechnol 42, 597–607 (2024).

