## Supplemental Figures for "Attenuating payload potency expands the therapeutic window of immunotoxins"

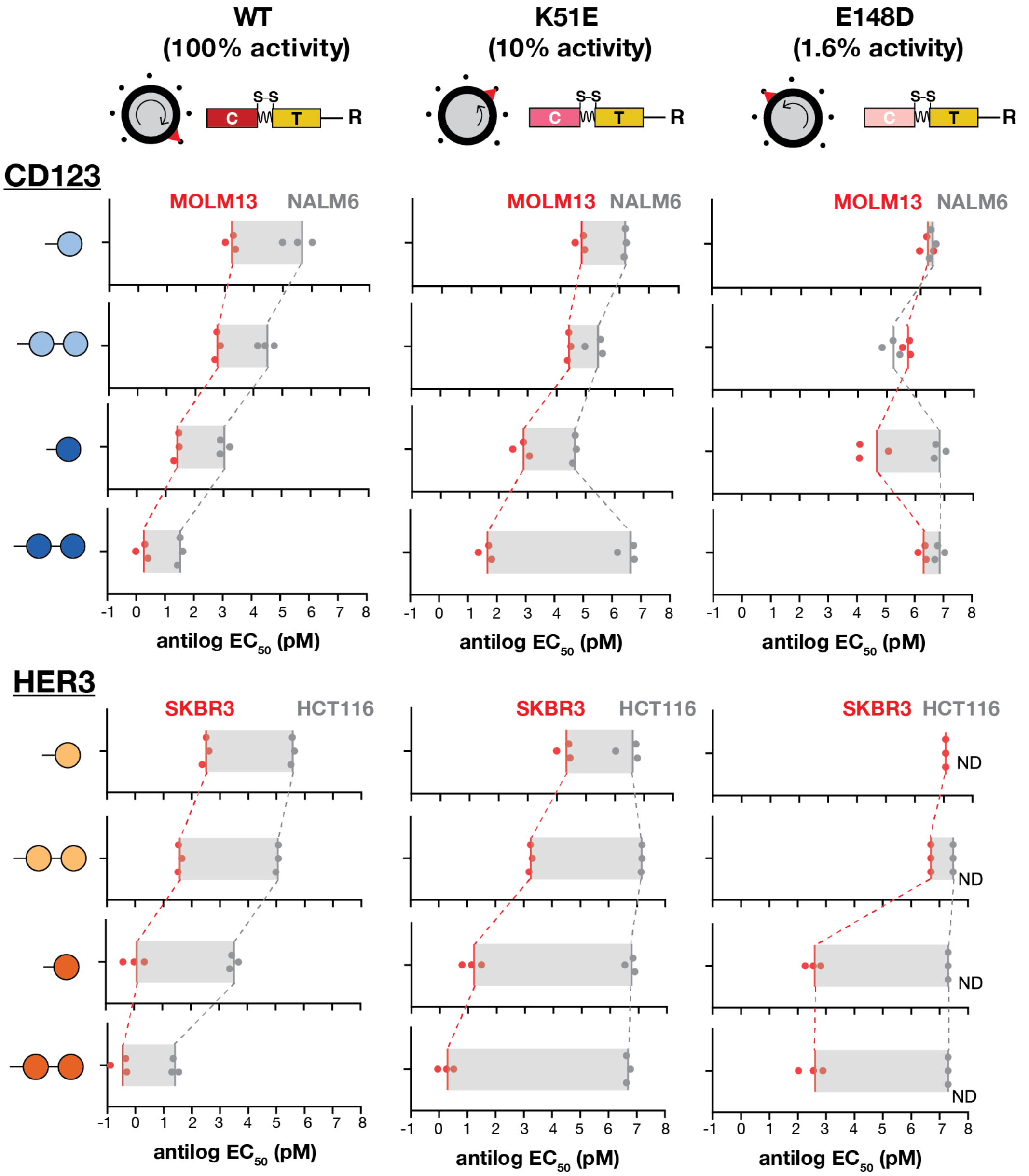


**Supplementary Figure 1. The effects of catalytic activity tuning on CD123- and HER3-targeted ITs.** CD123- and HER3-targeted ITs with WT DT-C (100% activity), K51E (around 10% activity) or E148D (around 1.6% activity) were evaluated across four RBD formats on either receptor-high and receptor-low cells. CD123-targeted ITs were tested in CD123-high MOLM13 cells (red) and CD123-low NALM6 cells (gray), whereas HER3-targeted ITs were tested in HER3-high SK-BR-3 cells (red) and HER3-low HCT116 cells (gray). Each point represents EC50 values from independent experiments, and the edge of the gray bar indicates the EC50 means (n=3, biological replicates). Gray bars indicate the magnitude of therapeutic window. ND: EC50 not determined because 50% cytotoxicity was not reached.

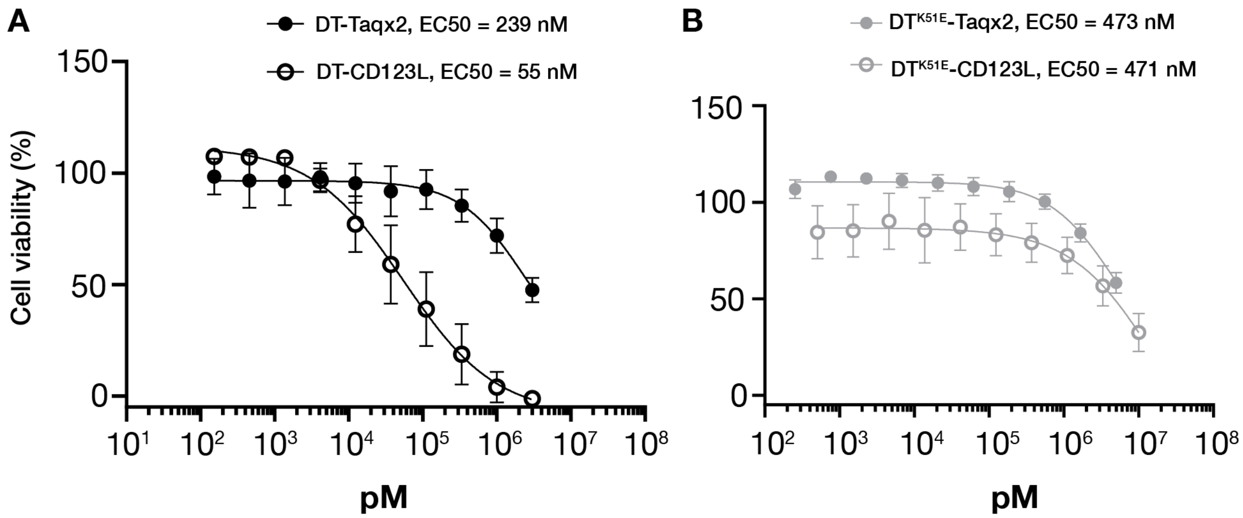


**Supplementary Figure 2. Activity tuning reduces receptor-mediated cytotoxicity in CD123-low cells.** WT (A) or K51E (B) CD123-targeted and untargeted ITs were assessed on CD123-low cells. Cell viability was measured 72 hours post treatment. n=3, biological replicates.

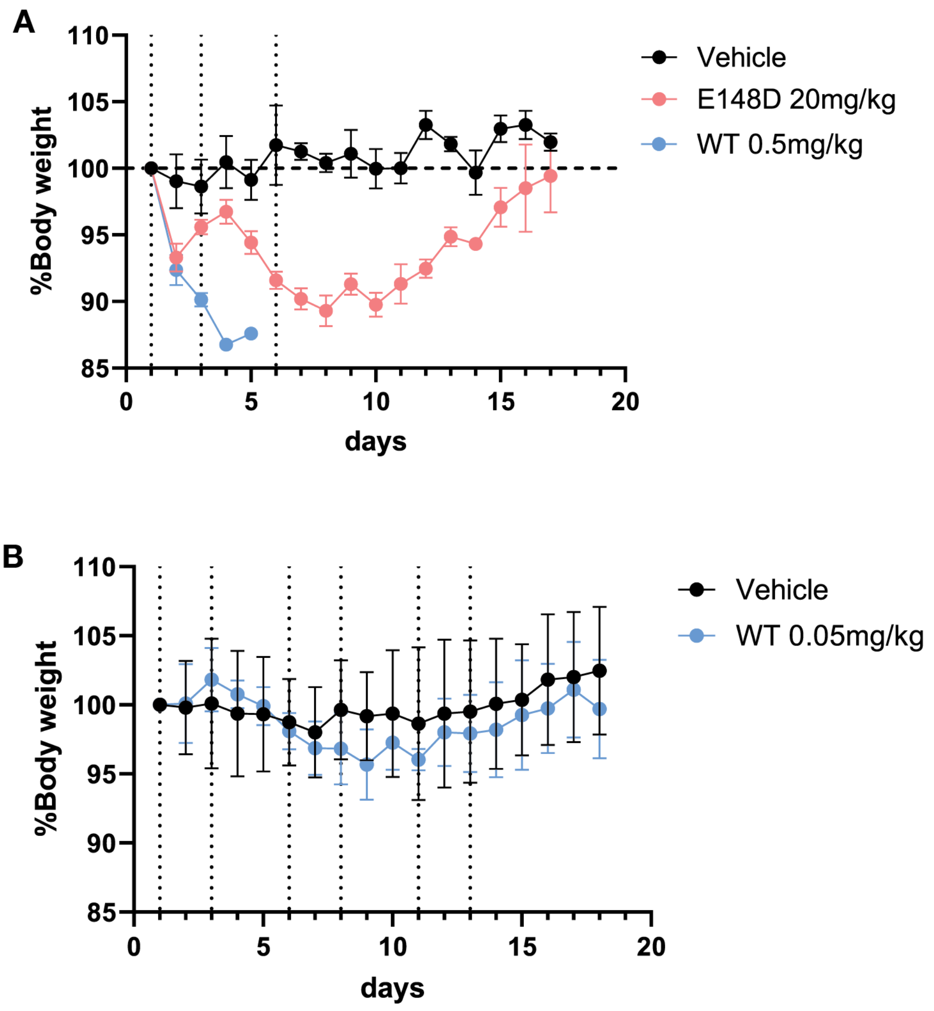


**Supplementary Figure 3. *In vivo* maximal tolerated doses of HER2-targeted ITs.** Body weight of NOD SCID mice was measured following the treatment of vehicle, DT-HER2$_H$x2-ABD or DT$^{E148D}$-HER2$_H$x2-ABD. (A). Mice were treated with vehicle, 20mg/kg of DT$^{E148D}$-HER2$_H$x2-ABD or 0.5mg/kg of DT-HER2$_H$x2-ABD. (B). Mice were treated with vehicle or 0.05mg/kg of DT-HER2$_H$x2-ABD. n=3.
